# Mobile-CRISPRi as a tool for genetic manipulation in the intracellular pathogen *Piscirickettsia salmonis*

**DOI:** 10.1101/2025.08.05.668709

**Authors:** Javiera Ortiz-Severin, Paulette Geoffroy, Pamela Aravena, Christian Hodar, Daniel E. Palma, Mauricio González, Verónica Cambiazo

**Author notes:** Correspondence: Verónica Cambiazo. Javiera Ortiz-Severín and Paulette Geoffroy contributed equally to this work and share first authorship. Author order was determined on the basis of seniority.

## Abstract

*Piscirickettsia salmonis* is the causative agent of Salmonid Rickettsial Septicemia (SRS), the main bacterial disease affecting the salmon industry in Chile. In this work, we implemented a Mobile-CRISPRi system to generate gene silencing using a catalytically inactive dCas9 protein and an IPTG-inducible single-guide RNA (sgRNA). We demonstrate the efficacy of the CRISPRi system in *P. salmonis* by silencing an exogenous gene encoding green fluorescent protein (sfGFP), and the endogenous homolog of the *fur* gene, whose gene product regulates intracellular iron homeostasis in bacteria. The inducible expression of *dcas9* and the *sfGFP*-directed sgRNA caused a 98.7% decrease in fluorescence in the knockdown strain. This silencing system was effective in seven *P. salmonis* strains from both genogroups. Furthermore, the same system was used to construct *fur* knockdown strains. A 50-fold decrease in *fur* expression level was determined in these strains, when the expression of the *fur* gRNA was induced with IPTG. By RNA-seq we detected a significant increase in the expression of genes encoding the Fe^+2^ and Fe^+3^ acquisition systems and iron mobilization in the *fur1* knockdown, after IPTG induction. All the genes with over two-fold increased expression in the RNA-seq presented the Fur box consensus sequence in their regulatory region. The successful implementation of the Mobile-CRISPRi system in *P. salmonis* paves the way for systematic analysis of gene function in this pathogen We anticipate that these analyses will be very valuable in identifying genes involved in the mechanisms of pathogenesis of *P. salmonis*.

**Importance:** Salmon Rickettsial Septicemia (SRS) is an infectious disease caused by the marine bacterium *Piscirickettsia salmonis*. This Gamma-proteobacteria is a fastidious and facultative intracellular pathogen that has a nearly worldwide distribution, particularly impacting Chilean salmonid aquaculture. Its fastidious nature has made it hard to grow in labs, hindering research into its virulence and treatment, especially because of the lack of molecular techniques to study gene function. We show here the successful implementation of the Mobile-CRISPRi system for gene silencing. Significantly, we have adapted this technique for use with the marine pathogen *P. salmonis*, inserting exogenous genes into the bacterium’s chromosome to ensure their constitutive and inducible expression, and silencing both exogenous and endogenous gene expression. The Mobile-CRISPRi system was also used to study the iron regulator Fur, confirming Fur’s relevance to the iron metabolism in the pathogen.

## Introduction

The CRISPR-Cas system has been broadly utilized in a variety of species as a genome engineering tool (Xu & Li, 2020). A related method, CRISPR interference (CRISPRi), uses an inactive Cas9 protein (dCas9) that has lost nuclease activity. Once the single-guide RNA (sgRNA) and dCas9 complex is formed, it represses transcription by inhibiting the initiation or progression of RNA polymerase at specific target locus. This system yields loss-of-function phenotypes that are suitable for a systematic genetic analysis aiming to uncover gene phenotypes (Banta *et al*., 2020; Larson *et al*., 2013; Peters *et al*., 2019). More recently, an additional system named as Mobile-CRISPRi has been established, including a collection of modular vectors that enable gene knockdowns in diverse Gram-negative bacteria by integrating IPTG-inducible *dcas9* gene and sgRNA sequences into the genome using the Tn7 transposon system (Peters *et al*., 2019). Therefore, Mobile-CRISPRi has the advantage of facilitating the analysis of gene function in non-model species and strains (Jin *et al*., 2022). Although the CRISPRi technology has been primarily characterized in the model bacteria *Escherichia coli* and *Bacillus subtilis*, there has been a recent proliferation of studies that have demonstrated the extension of these tools to other non-model bacteria, including those from the genera Mycobacterium and Streptomyces, the phyla Actinomycetota, Cyanobacteriota, Bacillota, and a wide variety of Pseudomonadota (Call & Andrews, 2022).

The objective of this study was to establish a stable and effective Mobile-CRISPRi induced knockdown for *Piscirickettsia salmonis*, a fastidious facultative intracellular bacterium that is the causal agent of Salmonid Rickettsial Septicemia (SRS), the main bacterial disease affecting the salmon industry in Chile (Sernapesca, 2024). *P. salmonis* has a nearly global distribution, being isolated in other important salmon-farming countries (Brocklebank, 1992; Olsen *et al*., 1997; Rodger & Drinan, 1993), where it is regarded as a potential emerging pathogen (Krkosek *et al*., 2024; Long & Jones, 2021). It is acknowledged that the range of molecular tools available to regulate gene expression in *P. salmonis* is limited, and that there is a clear constraint on our ability to interrogate the function of essential and conditionally essential genes.

In this study, using a Mobile-CRISPRi variant containing a constitutively expressed sfGFP reporter and an sgRNA targeting sfGFP (Peters *et al*., 2019) we tested the Mobile-CRISPRi system in different strains of *P. salmonis*, encompassing the two genogroups, LF89-like and EM90-like, previously described for this bacterium (Aravena *et al*., 2020). Afterwards, we used this system to study the function of the Ferric uptake regulator (Fur) in *P. salmonis*, which has been proven in other bacteria to reversibly bind Fe^2+^ and repress the expression of iron acquisition genes, allowing their expression only when the level of free intracellular iron is low (Liu *et al*., 2007). Fur has also been implicated in the regulation of virulence genes, stress resistance, toxin production and biofilm formation (Carpenter *et al*., 2009), and throughout small RNAs such as *rhyB*, or *IsrR*, Fur has been shown to indirectly regulate process such as the TCA cycle, and the response to oxidative stress (Coronel-Tellez *et al*., 2022; Massé *et al*., 2005; Massé & Gottesman, 2002; Troxell & Hassan, 2013). Due to its importance in virulence, the Fur regulon has been studied in several model bacterial species (Hall & Foster, 1996; Hickey & Cianciotto, 1994; Mey *et al*., 2005; Seo *et al*., 2014; Stojiljkovic *et al*., 1994). Iron metabolism is especially important in intracellular pathogens, such as *P. salmonis*, because iron acquisition from the host is essential for intracellular growth and survival, and thus, iron starvation becomes a virulence signal in these pathogens (Byrd & Horwitz, 1991; Collins, 2003). In a previous study, we predicted a set of iron acquisition genes in *P. salmonis* whose increased expression under iron-starved conditions, and the presence of putative Fur boxes upstream, suggested that most of these genes were part of the Fur regulon (Pulgar *et al*., 2015). Moreover, during *P. salmonis* infection in a macrophage-like cell line, metal acquisition proteins were one of the most highly expressed families of virulence factors (Ortiz-Severín *et al*., 2020). Here, we combined CRISPRi-based knockdown of fur with RNA sequencing to further characterize the Fur regulon of *P. salmonis*. Finally, we show that Mobile-CRISPRi is effective in *P. salmonis*, indicating the applicability of this system for gene functional analysis in this pathogen.

## Materials and methods

### 1. Bacterial strains and growth conditions

Oligonucleotides used in this study for PCR, qPCR and cloning can be found in Table S1. All bacterial strains and plasmids are listed in Tables S2 and S3, respectively. *P. salmonis* strains were routinely grown at 18°C with agitation, on Nutrient *Piscirickettsia* Broth (NPB; 30 g/L Tryptic Soy Broth, 256.6 mM NaCl, 8.25 mM L-cysteine, 37 µM FeCl_3_, and 2.5% of inactivated Fetal Bovine Serum (FBS). Bacteria were recovered from Nutrient *Piscirickettsia* Agar plates (NPA, NPB supplemented with 5% agar) and used to inoculate NPB. Cultures were incubated in a shaking incubator at 180 rpm and 18°C for 36 hours (exponentially growing bacteria) or 96 hours (stationary state bacteria) before harvesting. Purity of *P. salmonis* cultures was tested using a PCR-RFLP assay, as described in Mandakovic *et al*. (Mandakovic *et al*., 2016). *E. coli* strains were routinely grown in Luria-Bertani (LB) broth at 37°C with shaking. When required, the appropriate media were supplemented with gentamycin (30 μg/mL), ampicillin (100 μg/mL), or isopropyl β-D-1-thiogalactopyranoside (IPTG) at 100 μM unless otherwise noted.

### 2. Transformation of *E. coli* SM10

pJMP2754, pJMP2774 and pTNS3 plasmid DNA were purified from *E. coli* BW25141 and *E. coli* Pir1 hosts using the EZNA Plasmid Mini Kit II (Omega Bio-tek) following manufacturer’s instructions, quantified using Qubit Fluorometric Quantitation System (Life Technologies) and 50 to 75 ng of DNA were used to transform chemically competent *E. coli* SM10 cells following standard protocols. Transformed cells were plated on LB agar plates with gentamycin (30 μg/mL) or ampicillin (100 μg/mL) according to plasmid resistance.

### 3. Plasmid conjugation into *P. salmonis*

The Mobile-CRISPRi system was transferred to *P. salmonis* by tri-parental conjugation using the following *E. coli* donor strains: (1) SM10 strain containing the plasmid of interest (pJMP2754, pJMP2774, pJMP2782:sgRNA-*fur1*, or pJMP2782:sgRNA-*fur2*) and (2) SM10 strain containing pTNS3 encoding the Tn7 transposase. Before plasmid conjugation, a single colony of each *E. coli* strains was inoculated in 3 mL of LB broth with the appropriate antibiotic, and grown overnight (ON) at 37°C. The ON cultures were diluted to an optical density at 600 nm (OD_600_) of 0.01 in 5 mL of NPB medium supplemented with the appropriate antibiotic and grown at 20°C with shaking at 180 rpm for three days. *P. salmonis* recipient strain was also grown to stationary phase in NBP at 20°C with shaking for four days. One milliliter of donor strains culture was centrifuged at 6000 × g for 3 minutes at 4°C, washed three times with NPB, resuspended in 1 mL of NPB and mixed with the recipient strain in a ratio of 1:1:1 to a final volume of 300 µl. The mix was applied on a 0.45 µm S-Pak membrane filter (Merck) on NPA plates and incubated at 18°C for 40 hours. At the end of the incubation period, the filter was placed in a 50 mL conical tube containing 10 mL NPB and vortexed before 200 µL aliquots were poured on NPA plates supplemented with gentamycin (30 µg/ml) and trimethoprim (25 µg/ml). Plates were incubated at 18°C for two to three weeks before transconjugants were harvested. The matings were carried out in triplicate, using three independent cultures of each donor and recipient strain. At least ten colonies per conjugation event were picked up and inoculated on a 96 well plate with 200 µL NPB supplemented with gentamycin and trimethoprim and grown for 4 days at 20°C with shaking at 180 rpm. Grown cultures were checked by Gram-staining and PCR-RFLP (Mandakovic *et al*., 2016) for purity. The presence of plasmid insertion in *P. salmonis* genome was further confirmed by PCR with the Tn7R primer that hybridizes with the Tn7R sequence in the plasmid and glmsps2 primer that hybridizes to the 3’-end of *glmS* gene in *P. salmonis* chromosome (Table S1). Positive strains were grown in NPA plus gentamicin and trimethoprim for maintenance and downstream assays. Transconjugants were stored frozen in glycerol at −80°C.

### 4. Conjugative efficiency

The viable number of *P. salmonis* transconjugants was estimated by the Most Probable Number (MPN) method (Sutton, 2010). For that, 20 µL of the conjugation mixture per replicate (N = 5 independent matings) was inoculated into 180 µl of NPB supplemented with trimethoprim or with trimethoprim and gentamicin in 96-well plates, and the results were obtained from 12 ten-fold serial dilutions with 3 replicates for each dilution. The plates were incubated for 8-12 days at 20°C and the U.S. Environmental Protection Agency MPN calculator version 2.0 was used to generate the MPN counts for each sample (U.S. Environmental Protection Agency, 2013). Conjugative efficiency was calculated as the ratio of viable transconjugants grown in NPB supplemented with gentamycin plus trimethoprim (transconjugants), to viable *P. salmonis* cells grown in NPB plus trimethoprim (receptors).

### 5. Plasmid stability assay

Six isolates harboring Tn7 element from plasmids pJMP2754 or pJMP2774 were grown for 96 hours at 18°C in NPB supplemented with gentamycin and trimethoprim, then, 700 µL of each culture were centrifuged at 6000 × g for 3 min at 4°C and washed twice with NPB to remove the residual antibiotics. Washed cells were inoculated to a final OD_600_ = 0.01 in 700 µL NPB in a 48-well plate without antibiotics and grown for 96 hours at 18°C. The procedure of dilution and growth was repeated a total of 9 times (approximately 12 generations per growth experiment, a total of 132 generations). Then, retention of the Tn7 element was verified by PCR with primers Tn7R/glmsps2 and sfGFP fluorescence.

### 6. sfGFP fluorescence measurements

Transconjugant *P. salmonis* strains were grown for 40 h (mid-exponential phase) in NPB supplemented with gentamicin and trimethoprim. Then, 1 mL was centrifuged for 3 min at 6,000 × *g* at 4°C and resuspended in 1 mL of fresh NPB. After resuspension, 200 μL were directly added into sterile black 96-well plates or diluted 1:10, 1:100 or 1:1000 for fluorescence measurements (excitation 475 nm and emission 500-550 nm) in a GloMax Explorer Multimode Microplate Reader (Promega). Additionally, fluorescence of pJMP2754:*sfGFP* transconjugants (CGR02 *sfGFP*(+) strain) was detected using confocal microscopy. For this, 10 µL of stationary phase cultures were transferred to a 1% low melting point agarose pad (Lonza) and imaged. Expression of *sfGFP* was visualized using a C2+ Confocal inverted microscope (Nikon) and a 60 × objective, images were acquired using the NIS-elements program (Nikon).

### 7. Estimation of knockdown efficiency

Six independent transconjugant clones carrying Tn7 elements from pJMP2774 or pJMP2754 were grown to stationary phase and then inoculated in 24-well plates to a final OD_600_ = 0.01 in 1.2 mL of NPB, supplemented or not with 10 µM, 100 µM or 1000 µM of the inducer IPTG. Plates were incubated for 20, 44, 68 or 92 h at 18°C with shaking, and cultures were collected at each time point to quantify sfGFP fluorescence in a microplate reader as described above.

### 8. Growth curve analysis

For growth curve assays, stationary state bacteria (OD_600_ = 1.1-1.3) were used as inoculum. 48-well plates were inoculated to a final OD_600_ = 0.01 in 700 µl NPB broth and incubated at 18°C with shaking at 180 rpm. Every growth curve experiment was performed with three technical replicates and 11 biological replicates. Bacterial growth was assessed periodically by measuring the OD_600_ for five days in an Infinite 200 PRO NanoQuant (Tecan, Switzerland). Growth curve parameters were calculated based on raw growth data using the R package Growthcurver (Sprouffske & Wagner, 2016). Growth parameters were obtained for each biological replicate, and data were then processed in Excel to compare replicates separately. GraphPad Prism version 8.0.1 was used for graphical presentation and statistical analysis of the results.

### 9. Alignment of *attTn7* box in different *P. salmonis* strains

The 92 available RefSeq genomes for *P. salmonis* were downloaded from the NCBI website (https://www.ncbi.nlm.nih.gov/datasets/genome/?taxon=1238 accessed on July 23^th^, 2025). For each genome, the sequence of the *glmS* CDS plus 30 nucleotides downstream were extracted using BEDTools v2. 31.1 (Quinlan & Hall, 2010) and aligned with MAFFT v7.526 (Katoh & Standley, 2013). Finally, a sequence logo of the TnsD binding region (*attTn7* box) was generated using WebLogo (Crooks *et al*., 2004).

### 10. sgRNA design

The sgRNA spacers were designed as described in Banta *et al*. (https://github.com/ryandward/sgrna_design) (Banta *et al*., 2020). Briefly, 20 nucleotide sequences were selected using the following criteria: next to a NGG protospacer adjacent motif (PAM), targeting the non-template strand towards the 5′ end of the gene, and having maximum specificity when aligned to the target genome. Alignment with *P. salmonis* CGR02 genome (NCBI RefSeq assembly GCF_001534725.1) was performed using Bowtie (Langmead *et al*., 2009). Two oligonucleotides, forward and reverse, were designed to obtain the selected sgRNA when annealed. Each oligonucleotide has an additional 4 bp sequence at the 5’ end that is complementary with the *BsaI* sticky ends that are generated in pJMP2782 plasmid enabling ligation into the plasmid. Two sgRNA spacers targeting *fur* gene were designed and named sgRNA-*fur1* and sgRNA-*fur2*.

### 11. Vector construction

Plasmid DNA was extracted from 15 mL *E. coli* BW25141 pJMP2782 culture using EZNA Plasmid Mini Kit I (Omega Bio-tek) following manufacturer’s instructions, digested using the *BsaI* restriction endonuclease (New England Biolabs) for 10 hours at 37°C, purified using Zymo DNA clean and concentrator-5 kit, and quantified using Qubit Fluorometric Quantitation System (Life Technologies). The oligonucleotides were annealed by incubating them at 95°C for 5 minutes and allowing them to cool to room temperature. This final solution was diluted 40 times. Ligations of the dsDNA oligos to the digested pJMP2782 plasmid were performed with T4 DNA ligase (NEB) in a 10 μL reaction volume with 50 ng of digested pJMP2782 and 100 μM of each oligo. The reaction was incubated for 16 hours at 16 °C followed by heat-inactivation for 10 minutes at 65 °C. Reactions were transformed into chemically competent *E. coli* SM10 cells as previously described, after growth in selective media, *P. salmonis* transconjugants (sgRNA-*fur1* and sgRNA-*fur2*) were checked by PCR as mentioned above.

### 12. RNA purification

*P. salmonis* strains sgRNA-*fur1* and sgRNA-*fur2* were grown for 38 hours (mid-exponential phase) at 18°C in NPB supplemented or not with IPTG 100 µM and then were collected by centrifugation 8000 × g for 10 min at 4°C, supernatants were discarded and the pellets were suspended in 750 µL of TRIzol (Thermo Fisher Scientific) and total RNA extraction was carried out according to manufacturer’s instructions. Four micrograms of RNA were treated with Turbo DNA-free Kit (Invitrogen) according to standard protocols. Purified RNA was resuspended in 50 µL of nuclease free water and quantified using a Qubit Fluorometric Quantitation System (Life Technologies). RNA quality was assessed using the 2200 TapeStation Bioanalyzer (Agilent Technologies).

### 13. Quantitative real-time PCR (qPCR)assays

qPCR reactions were carried out in an AriaMx Pro thermal cycler (Agilent Technologies). cDNAs were synthesized from 1 µg of DNA-free RNA using the High-Capacity RNA-to-cDNA Kit (Applied Biosystems) according to manufactureŕs instructions. cDNAs were diluted to 5 ng and used as template for qPCR, with primers designed against the genes of interest (see Table S1). PCR conditions were 95°C for 3 min followed by 95°C for 3 s, 62–62.5°C for 10 s and 72°C for 12 s for a total of 35 cycles. Melting curves (0.5°C steps between 65–95°C) ensured that a single product was amplified in each reaction. To determine relative expression levels of genes, the method described by Pfaffl (Pfaffl, 2001) was employed, using gene *recF* as an internal reference gene (housekeeping). At least 3 biological replicates were analyzed, and PCR efficiencies were determined by linear regression analysis performed directly on the sample data using LinRegPCR (Ramakers *et al*., 2003).

### 14. RNA sequencing and analysis

Three libraries for control (uninduced sgRNA-*fur1* strain) and three for IPTG-induced sgRNA-*fur1* strain were generated at SeqCenter (Pittsburgh, PA), using the TruSeq Stranded Total RNA Library Prep (Illumina), with bacterial rRNA removal. The libraries were paired-end sequenced on a NovaSeq X Plus instrument (Illumina) with read lengths of 150 bp. The RNA-seq raw data quality was examined using FastQC (S. Andrews, 2010) to detect low quality reads and Illumina adapters, which were subsequently eliminated using BBduk software (Bushnell, 2014). The filtered reads were mapped against the *P. salmonis* CGR02 genome (GenBank accession N° GCA_001534725.1) using the STAR v2.7.10b RNA-Seq alignment tool. Reads that failed to map to any gene or mapped to multiple genes were removed before expression analysis. Gene quantification was generated using the quantMode option available at STAR program. To identify the differentially expressed genes (DEGs), the gene counts matrices were previously filtered by removing genes with counts in less than two of three replicates by condition. Finally, low expressed genes were discarded using the strategy described in (Chen *et al*., 2016). RNA-seq data can be accessed at NCBI, BioProject PRJNA1300335.

DESeq2 v1.46.0 package (Love *et al*., 2014) was used to detect DEGs. First, raw count data was used to calculate normalization factors to correct for variations in library depth. Next, gene-specific dispersion values were estimated and refined through shrinkage of gene-wise dispersion. Finally, we fit a negative binomial model to each gene followed by a differential expression test using the Wald significance test. All these calculations were carried out using R software v.4.3.1. Plots were constructed using the R package ggplot2 (v.3.4.4) (Wickham, 2016). For functional analysis of differentially expressed genes, a compilation of gene ontology (GO) terms for *P. salmonis* CGR02, was generated using eggNOG mapper v2.1.9 (Cantalapiedra *et al*., 2021), and an org.db package was constructed using the R package AnnotationForge v 1.42.2. Then, GO enrichment analysis was carried out using enrichGO function from R package clusterProfiler (v.4.8.1) (Wu *et al*., 2021). Additionally, homologs of the Virulence Factor Database (VFDB) proteins (downloaded on 21st April 2023) were identified using DIAMOND v2.0.15 (Buchfink *et al*., 2021). *P. salmonis* CGR02 genome was also screened for the genes of iron metabolism with the bioinformatics tool FeGenie (Garber *et al*., 2020).

### 15. Prediction of Fur-binding sites

For the identification of potential Fur-binding sites (Fur boxes) in the *P. salmonis* genome, 774 experimentally validated transcription factor binding sites were downloaded from the CollectTF database (Kiliç *et al*., 2014). The sequences were converted to the MEME motif format (Bailey *et al*., 2015) and scanned against the *P. salmonis* genome using the command line FIMO tool (Grant *et al*., 2011). The parameters of the scanning were first tested using *E. coli* genome and selected when the 95% of the already described Fur-binding sites on *E. coli* genome were identified (Seo *et al*., 2014). The Markov background DNA model for *P. salmonis* genome was constructed using fasta-get-markov script from MEME suite. Finally, binding site searching was carried out with the following parameters: -alpha 0.1 -oc furBkg0Motif2024 --max-strand -- bgfile psal.mk0.bgk --parse-genomic-coord --thresh 0.0001. Then, sequence logos were created using R package ggseqlogo v0.2 (Wagih, 2017).

### 16. Quantification of intracellular iron and proteins

Exponentially growing *P. salmonis* cells (1 mL) were collected by centrifugation at 8000 × g for 10 min at 4°C. The supernatant was discarded, and the pellet was subsequently washed with PBS 1X, 50 mM glycine, 1 mM EDTA and milliQ water. Following centrifugation at 10,000 × g for 10 min at 4°C, the resulting pellet was resuspended in 125 µL of 65% nitric acid (Merck) and incubated at 62°C overnight to fully dissolve the pellet. Total reflection X-ray fluorescence spectroscopy (TXRF) with gallium as the internal standard was performed using a Bruker S2 PICOFOX, as described by González *et al*. (González *et al*., 1999).

To quantify the protein content of the bacteria, 1 mL of the samples was centrifuged at 8000 × g for 10 min at 4°C. The supernatant was then discarded, and the pellet resuspended in PBS containing a protease inhibitor (Sigma-Aldrich). The samples were sonicated in two 30-second pulses, incubating on ice between each pulse, then they were centrifuged again at 10,000 × g for 10 min at 4°C. Quantification was performed using the Qubit Protein Assay Kit (Thermo Fisher), following the supplier’s instruction. The concentration of each sample was then determined by interpolation using a calibration curve constructed with 3 internal standards. GraphPad Prism (GraphPad Software, La Jolla, CA, United States) was used for graphical presentation and statistical analysis of data.

## Results

### Mobile-CRISPRi functions in *P. salmonis* strains

In order to test the functionality of the Mobile-CRISPRi platform in *P. salmonis*, we used the plasmids pJMP2754 and pJMP2774, which were designed by Banta *et al* (Banta *et al*., 2020). The pJMP2754 plasmid contains the “test” version of the Mobile-CRISPRi, which consists of a superfolder green fluorescent protein gene (*sfGFP*) with constitutive expression and an IPTG-inducible *dcas9* gene, but lacks the sgRNA. We used this plasmid to produce a control strain named *sfGFP*(+). We used pJMP2774, which contains an IPTG-inducible sgRNA (designated as gmc6) targeting the *sfGFP* gene, to generate the strain sgRNA-*sfGFP*. A schematic representation of the pJMP2774 Mobile-CRISPRi and the predicted insertion site in the genome of *P. salmonis* CGR02 are shown in Fig. 1A. We examined the conservation of the Tn7 transposon site in the genomes of 92 strains of *P. salmonis*, including representatives of the two genogroups described for this species. Previous works have shown that the TnsD protein recognizes and binds to a DNA sequence called *attTn7* box near the end of the highly conserved *glmS* gene and subsequently inserts the transposon downstream of the GlmS open reading frame. Consistently, our analysis of the *attTn7* binding site of *P. salmonis* showed that it is highly conserved in the 92 sequences of *P. salmonis* strains (Fig S1).

**Fig 1.**
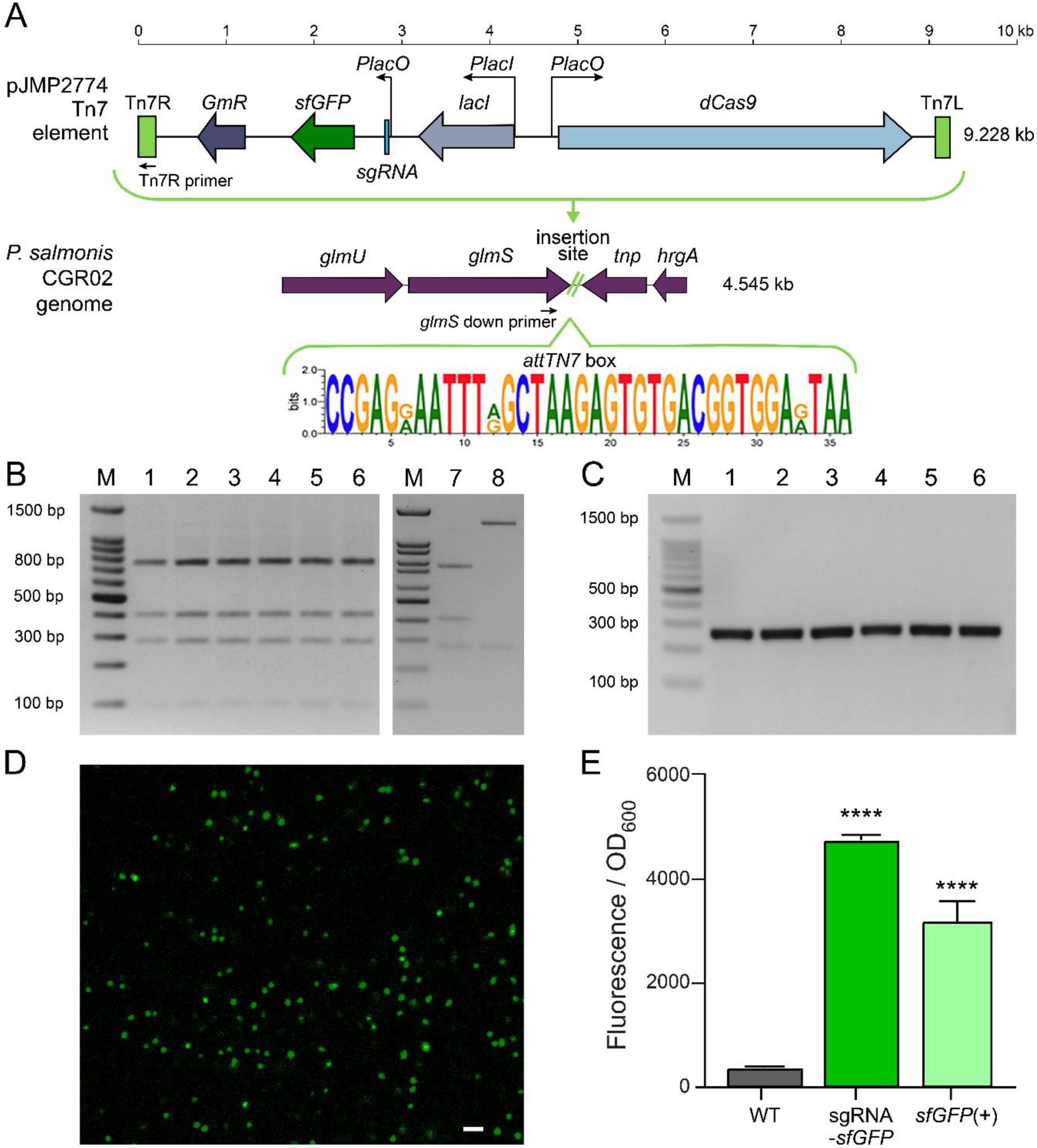
Tn7 integration and functional demonstration of Mobile-CRIPSPRi system in *P. salmonis* CGR02. (A) Model depicting Tn7 insertion of Mobile-CRISPRi in the genomic context of *P. salmonis* CGR02, and the consensus *attTn7* box (TnsD recognition sequence). The Tn7 element in pJMP2774 contains Tn7R, Tn7L, Gentamycin resistance cassette (*GmR*), and foreign genes *dcas9*, *sfGFP* and the gmc6 sgRNA targeting the *sfGFP* gene. The predicted alignment sites of primers Tn7R and glmsps2 down for the experimental confirmation of the insertion are also shown. (B) PCR-RFLP digestion pattern (16S rDNA, amplification using primers 27F/1492R) for the *sfGFP*(+) and sgRNA-*sfGFP* transconjugants*. P. salmonis* CGR02 and *E. coli* SM10 are included as controls of PCR-RFLP patterns (lanes 7 and 8, respectively). (C) PCR confirmation of Mobile-CRISPRi insertion using primers Tn7R/glmsps2 down shown in (A), with an expected fragment of ∼300bp. (B) and (C), lanes M indicate the molecular weight standards, lanes 1-3 different *P. salmonis* sgRNA-*sfGFP* transconjugants and lanes 4-6 show *sfGFP*(+) transconjugants. (D) Confocal microscopy showing the green fluorescence of CGR02 *sfGFP*(+). Bar = 10 µm. (E) The GFP fluorescence of the *P. salmonis* sgRNA-sfGFP and sfGFP(+) transconjugants harboring the Mobile-CRISPRi system, which was not induced to express dCas9 and the sgRNA, was compared with that of the wild-type strain (WT). Fluorescence was normalized according to the optical density (OD_600_) of each culture IPTG (unpaired t-test, **** p < 0.0001, n = 4 independent replicates).

Considering the fastidious nature of *P. salmonis*, the bacterium was routinely cultured in nutrient media (NPB or NPA) at 18°C. The conjugation experiments were also conducted in these conditions, which allowed the growth of both *P. salmonis* and *E. coli*. To decrease the chance of co-culturing both bacteria after conjugation, transconjugants were selected in NPA medium supplemented with gentamycin (to select for the insertion of the Mobile-CRIPSPRi module) and with trimethoprim, since *P. salmonis* is naturally resistant to trimethoprim and sensitive to gentamycin (Fig. S2; (Ortiz-Severín *et al*., 2021). In addition, PCR-RFLP was performed for every *P. salmonis* transconjugant to ensure its purity (no carry-over of *E. coli* cells, Fig. 1B). Afterwards, the correct insertion of the Mobile-CRIPSPRi module was confirmed by the amplification of the expected ∼300 bp sequence obtained by using Tn7R primer and the *glmS* down primer (Fig. 1A, Table S1) in the transconjugant clones (Fig. 1C). We measured Mobile-CRISPRi transfer quantifying the number of recipient cells (transconjugants) by determining the MPN on gentamycin + trimethoprim NPB as a fraction of total *P. salmonis* cells recuperated in trimethoprim supplemented NPB, *P. salmonis* CGR02 showed transfer efficiencies between 1.00 x 10^-1^ and 2.57 x 10^-1^ depending upon the experiment. Bright green fluorescence of *P. salmonis* transconjugants harboring the *sfGFP* gene (*sfGFP*(+)) was detected by confocal microscopy (Fig. 1D). Furthermore, in relation to the wild type (WT) strain, a significant increase in fluorescence was detected in the transconjugants strains harboring the Mobile-CRISPRi system without *dcas9* induction (Fig. 1E).

The inserted Mobile-CRIPSPRi module remained stable in the *P. salmonis* CGR02 genomic context. Following ten subcultures in the absence of antibiotic selection pressure, the transconjugants retained the inserted module. (Fig. 2A). Furthermore, following over 130 generations, the uninduced *P. salmonis* strains sgRNA-*sfGFP* and *sfGFP*(+) retained the ability to emit fluorescence (Fig. 2B).

**Fig 2.**
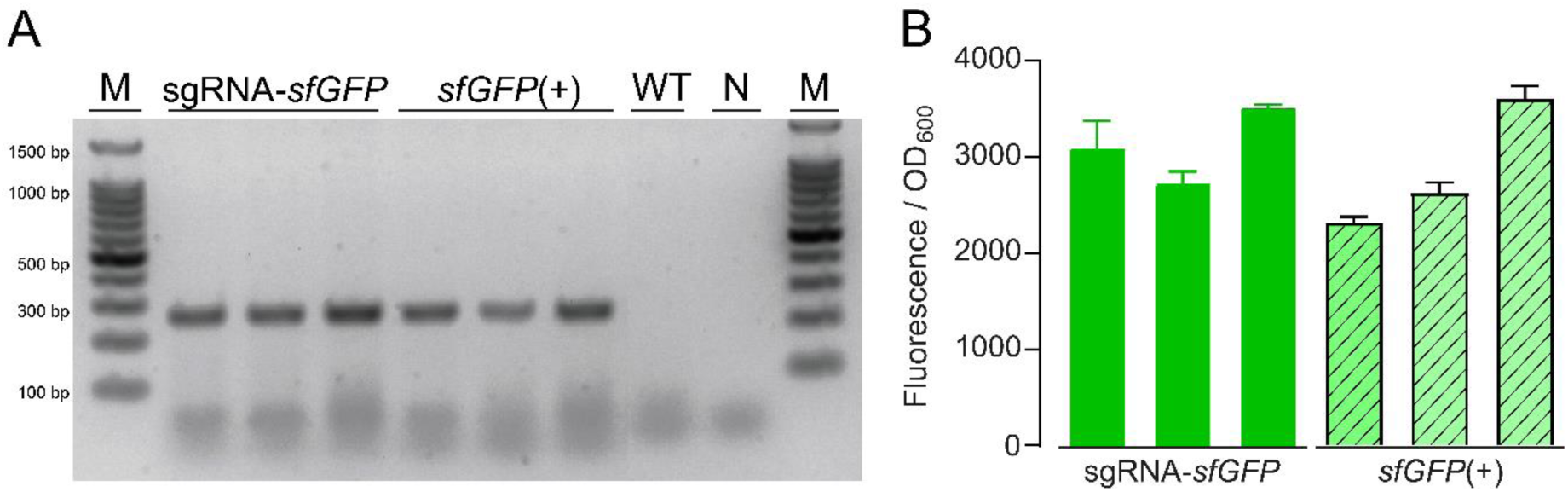
Stability of the Mobile-CRIPSPRi system in *P. salmonis.* (A) The gel electrophoresis shows the PCR product to confirm presence of the insertion using Tn7R/glmsps2-down primers for three different *P. salmonis* clones harboring the Mobile-CRISPRi systems (*sfGFP*(+) or sgRNA-*sfGFP*), the wild-type strain (WT) and a PCR-negative control (N). Lanes M show the molecular weight standard. (B) Relative fluorescence normalized by the optical density (OD_600_) of each culture, with three biological replicates and three technical replicates each. The graph shows three different clones of uninduced *P. salmonis* transconjugants.

To demonstrate the efficacy of the silencing system in the genetic context of *P. salmonis*, the expression of *dcas9* and sgRNA-*sfGFP* was induced using different concentrations of IPTG. As shown in Fig 3, the relative fluorescence decreased in the presence of the inducer. In subsequent experiments, 100 µM of IPTG was used, as it was the lowest concentration with the greatest effect, showing at 68 h a knockdown efficiency of 98.7%, similar to that of 1000 µM (98.7%), but significantly higher than that of 10 µM of IPTG (89.7%). Furthermore, the induction of *dcas9* in conjunction with the *sfGFP* targeting sgRNA, or the increasing concentrations of IPTG, did not result in any alterations to the growth of the *P. salmonis* transconjugants in NPB (Fig. 3).

**Fig 3.**
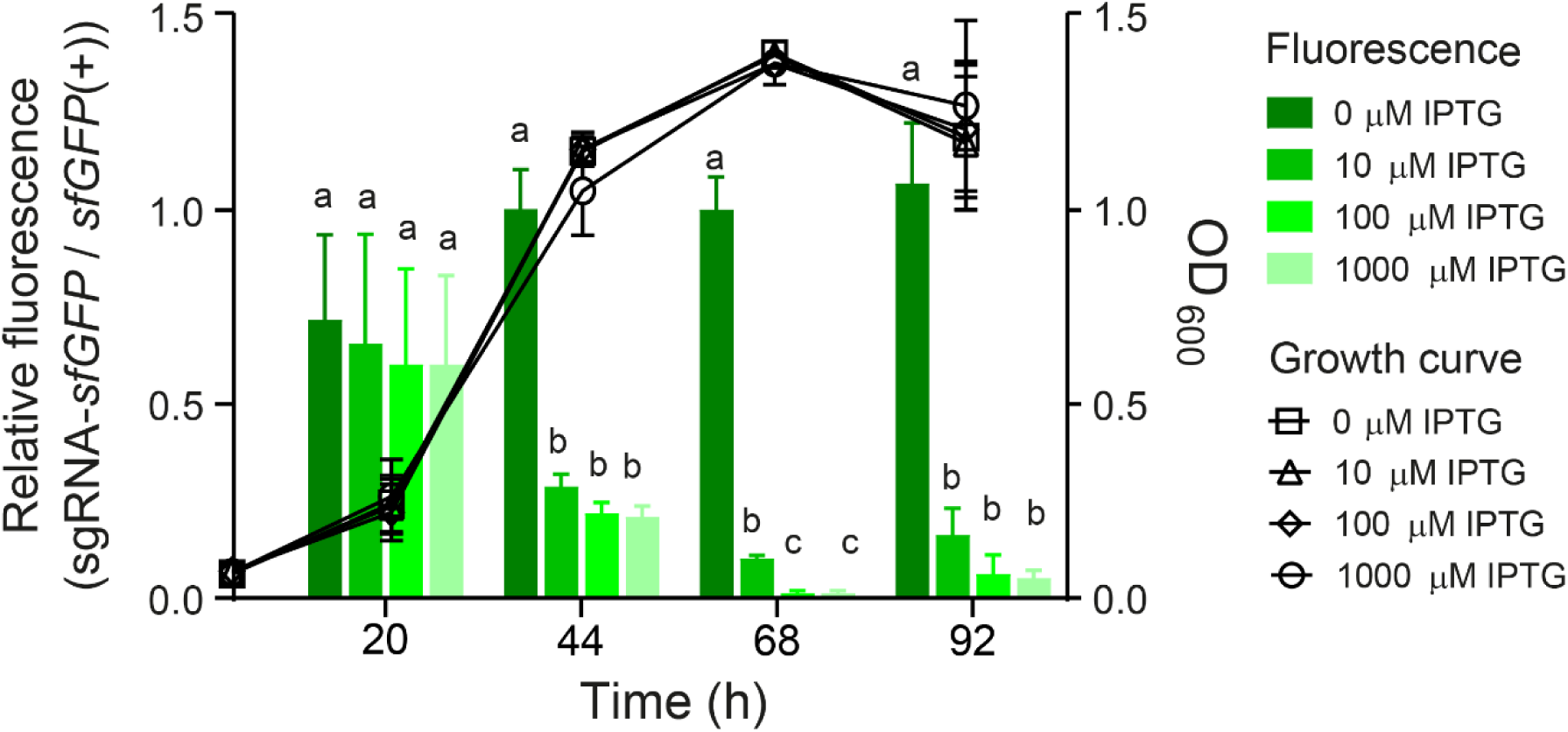
Silencing of *sfGFP* expression via Mobile-CRISPRi in *P. salmonis* CGR02. The green bars represent the relative fluorescence (fluorescence measurements normalized by OD_600_) of the inducible sgRNA-*sfGFP* strain, divided by the *sfGFP*(+) strain at different time points with increasing IPTG concentrations (0 µM, 10 µM, 100 µM or 1000 µM). Each data point is based on three biological replicates. The black symbols show the growth curve of *P. salmonis* harboring sgRNA-*sfGFP* with different concentrations of IPTG. Left Y axis shows relative fluorescence and right Y axis shows optical density (OD_600_). Letters over the bar-graphs indicate statistical differences (evaluated by one-way ANOVA for each time point and Tukey multiple comparisons post-test).

In order to provide further evidence for the broad application of the Mobile-CRISPRi silencing system in *P. salmonis*, the system was applied to two strains of genogroup EM90-like (*P. salmonis* 12201 and 8079) and four strains of genogroup LF89-like. Of these, two were isolated in Chile (*P. salmonis* LF-89 (ATCC VR-1361) and PSCGR01), one was a Norwegian strain (NVI5692), and one was a Canadian strain (NVI5892). All strains were transformed using the pJMP2774 plasmid (Table S3). As a result, all strains tested were able to conjugate with *E. coli* and all transconjugants contained the inserted Mobile-CRISPRi module. Moreover, all the strains were able to express fluorescence and to silence *sfGFP* expression in the presence of IPTG (Fig. 4).

**Fig 4.**
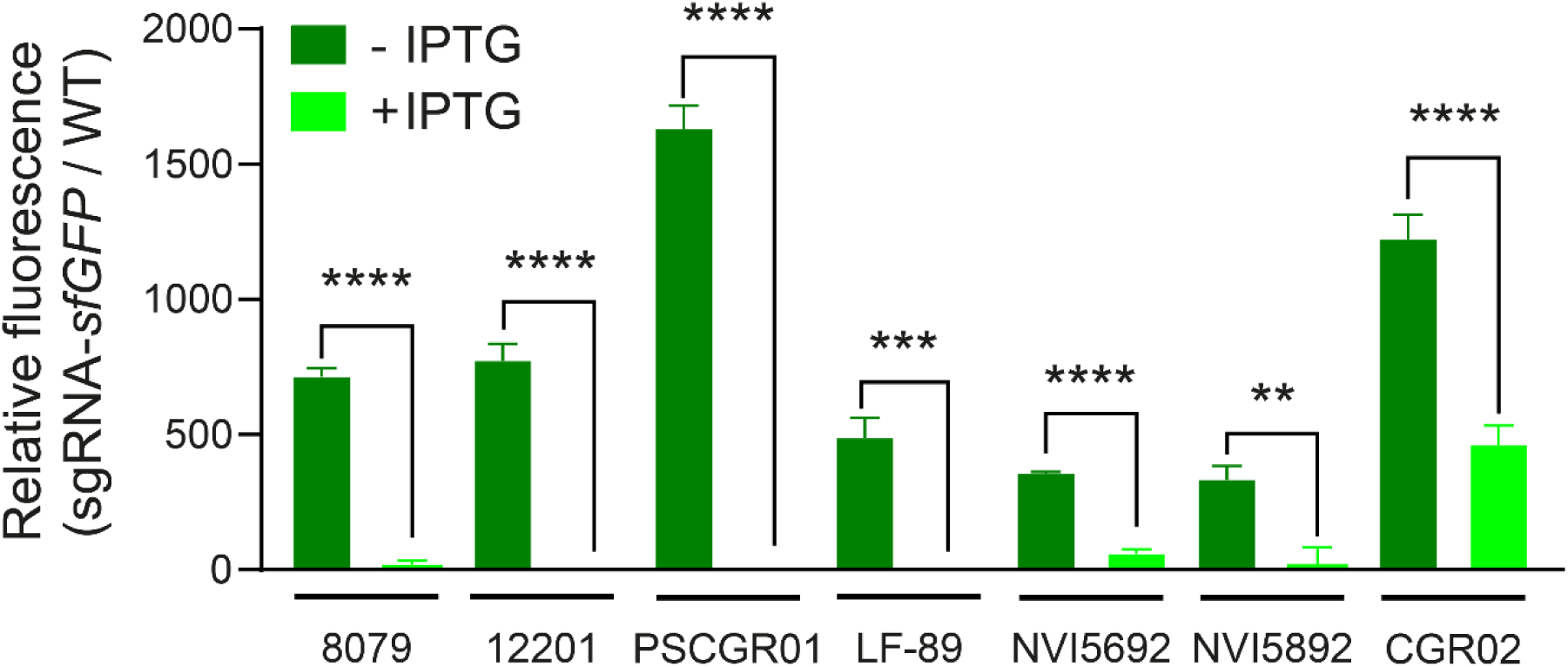
Fluorescence of different *P. salmonis* strains carrying the Tn7 element from the Mobile-CRISPRi silencing system. The green bars show the relative fluorescence of the different *P. salmonis* strains normalized by the optical density of the culture, the normalized by the fluorescence of the WT strain. Three replicates of each *P. salmonis* strain are shown, with or without supplementation of the inducer IPTG (100 µM). Statistical significance was evaluated using an unpaired t-test comparing each strain with its control without IPTG. (** p < 0.01, *** p < 0.001 **** p < 0.0001).

### Mobile-CRISPRi targets the *fur* gene

To further test the utility of the Mobile-CRISPRi system, two Mobile-CRISPRi variants were constructed containing sgRNAs targeting the *fur* gene of *P. salmonis*, encoding the ferric uptake regulator (Fur) protein, which is known to act as a Fe^2+-^dependent transcriptional repressor of bacterial iron acquisition genes (Escolar *et al*., 1999). Furthermore, this gene encodes a monocistronic mRNA, which makes the polar effect of CRISPRi unlikely. All the analyses of gene expression and growth curves of the *fur* knockdown RNA strains (sgRNA-*fur1* and sgRNA-*fur2*), induced and uninduced by IPTG, were carried out in the NPB medium supplemented with 400 μM FeCl_3_. This is an iron concentration that significantly increased the relative expression of *fur* measured by qPCR (Fig. 5A), without altering the growth curves obtained, which were indistinguishable from those of the WT strain. (Fig. 5B).

**Fig 5.**
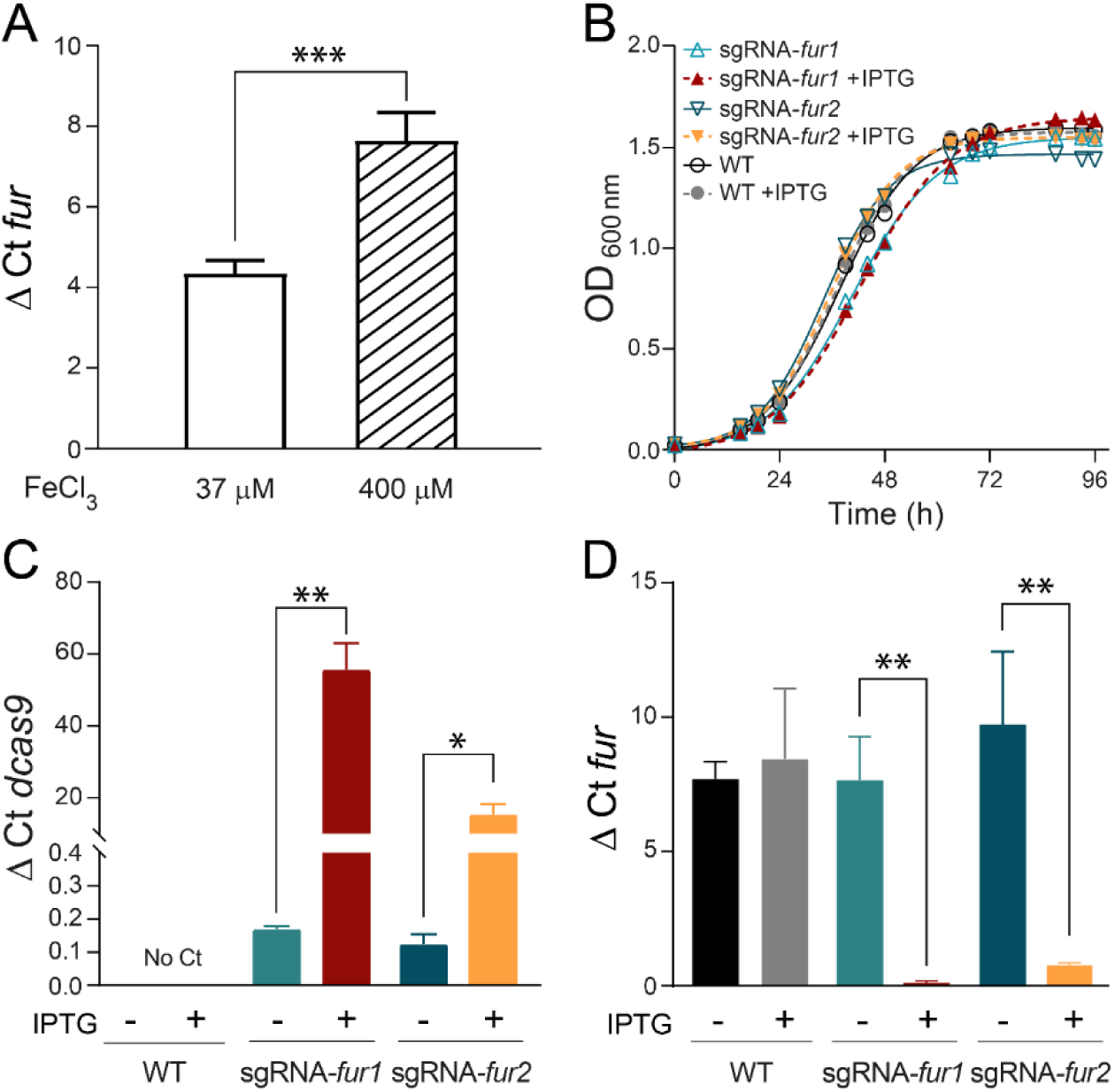
Silencing of *fur* expression via Mobile-CRISPRi in *P. salmonis* CGR02. (A) *fur* expression in the WT strain grown in culture media supplemented with 37 µM or 400 µM FeCl_3_. Expression levels (ΔCt) were normalized by the housekeeping gene *recF* (unpaired t-test, *** p < 0.001, n = 3-6). (B-D) Two sgRNA were designed to target *P. salmonis* endogenous *fur* and used to create the knockdown strains sgRNA-*fur1* and sgRNA-*fur2*. (B) Growth curves of *P. salmonis* WT, sgRNA-*fur1* and sgRNA-*fur2* strains with or without IPTG supplementation. (C-D) Quantification of gene expression in *P. salmonis* WT, sgRNA-*fur1* and sgRNA-*fur2* by qPCR is shown as ΔCt values (normalized by the expression of the housekeeping gene *recF*) for *dcas9* (C) and *fur* (D). Results are shown for each strain with or without supplementation of the inducer IPTG (unpaired t-test, * p < 0.05, ** p < 0.01, n = 3-6 independent replicates).

We quantified the expression of *dcas9* in sgRNA-*fur1* and sgRNA-*fur2* strains by qPCR in the absence or presence of IPTG induction. Basal levels of *dcas9* expression were detected in the uninduced strains, and a significant increase in *dcas9* expression was observed upon induction of sgRNA-*fur1* and sgRNA-*fur2* strains with IPTG (Fig. 5C). We then characterized Mobile-CRISPRi efficiency to silence the endogenous *fur* gene by qPCR. As expected, the levels of *fur* mRNA were significantly decreased in the presence of IPTG, both in sgRNA-*fur1* and sgRNA-*fur2* strains (Fig. 5D). IPTG induction of *dcas9* and sgRNA expression for 40 hours in sgRNA-*fur1* and sgRNA-*fur2* strains depleted *fur* mRNA by at least 95% of the levels measured in the corresponding non-induced or WT strains in the absence of IPTG (Fig. 5D). As demonstrated in Figure 5B, the induction of dCas9 and sgRNA expression at a previously defined concentration of 100 µM IPTG did not result in a significant growth defect, thereby ruling out potential toxic effects due to overproduction of dCas9, as observed in other bacteria (Rock *et al*., 2017; Zhang & Voigt, 2018). This result also indicates that the knockdown of *fur* does not have a detrimental effect on the viability of *P. salmonis* during growth in NBP.

### RNA-seq characterization of CRISPRi-mediated *fur* knockdown

Transcriptome analysis was performed on IPTG-induced and uninduced sgRNA-*fur1* strains under iron-replete condition to elucidate the global Fur-dependent regulation (Figure 6). The repressive regulatory effects exerted by Fur in wild-type bacteria under iron-replete conditions should be alleviated due to the knockdown of Fur in the IPTG-induced strain. RNA samples were prepared from three biological replicates of the exponentially growing sgRNA-*fur1* strain after two passages in NPB plus 400 μM FeCl3, with and without IPTG. In these conditions, the depletion of *fur* was found to be significantly higher than that observed after a single passage in broth supplemented with IPTG (see Fig. S3). The RNA-seq data were analyzed for accuracy (adjusted p-value < 0.05), revealing 219 differentially expressed genes. Filtering by fold change (log₂ FC ≥ 1, log₂ FC ≤ −1; see Table S4), we found that the expression level of 69 genes (45 upregulated and 24 downregulated) changed significantly between the IPTG-induced and uninduced sgRNA-*fur1* strains (see Fig. 6A).

**Figure 6.**
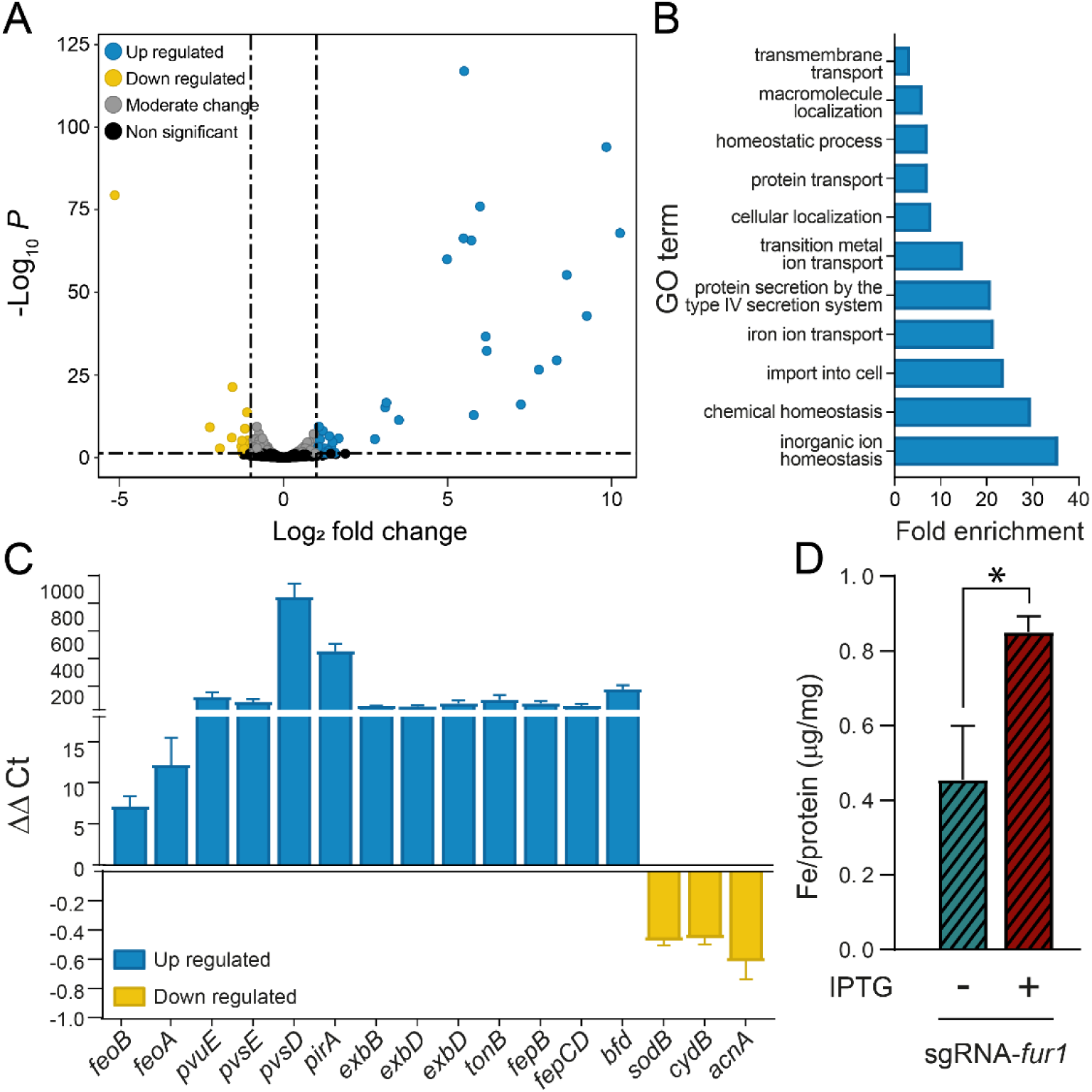
Comparative transcriptomic analysis of *fur*-knock down in *P. salmonis* CGR02. (A) Volcano plots from the DESeq2 analysis of differentially expressed genes between *P. salmonis* sgRNA-*fur1* knockdown strain and the isogenic strain without IPTG inducer (control). The blue dots (n = 45) represent significantly upregulated genes, the yellow dots (n = 24) represent significantly downregulated genes (adjusted p-value <0.05, |log_2_FC| > 1). Grey dots represent no significant change of expression. (B) GO enrichment analysis of upregulate genes. List of significantly overrepresented GO terms in the DEGs (log_2_ FC > 1) in the sgRNA-*fur1* knockdown strain. (C) RT-qPCR quantification of transcripts for the most up or downregulated genes identified in the RNA-seq analysis. Expression levels of *P. salmonis* genes are expressed as ΔΔCt values of the sgRNA-*fur1* knockdown strain normalized by the control, and *recF* as housekeeping gene. Increased gene expression relative to control bacteria is shown in blue, and decreased expression in yellow. (D) Intracellular iron concentration in sgRNA-*fur1* strain with (+) or without (-) the inducer IPTG (unpaired t-test, * p < 0.05, n = 3 independent replicates).

From Table S4, it becomes evident that the genes with the highest fold change were those upregulated in the IPTG-induced sgRNA-*fur1* strain. Moreover, the number of genes upregulated was higher (66%) than the genes downregulated (34%) in the knockdown strain. GO enrichment analysis indicated that the biological processes “inorganic ion homeostasis”, “import into cell”, “iron ion transport” and “secretion by the type IV secretion system” were enriched among the genes upregulated in the sgRNA-*fur1* strain (adj. p-value < 0.05, fold enrichment > 2; Fig. 6B), however, no enriched biological process were detected for downregulated genes. Among the most highly expressed genes in the induced sgRNA-fur1 strain (N = 18, adjusted p-value < 0.05, log₂ FC >2, Table 1), we identified genes encoding putative components of the FeO and FeP systems, which are involved in ferric and ferrous transport, respectively (Andrews *et al*., 2003). We also identified the *bfd* gene, which encodes a bacterioferritin-associated ferredoxin required for mobilizing iron stored in bacterioferritin (Yao *et al*., 2016). The highest fold inductions in the knockdown strain were observed for a gene encoding a putative TonB-dependent siderophore receptor (PirA), members of the TonB-ExbB-ExbD energy transduction apparatus (Ferguson & Deisenhofer, 2004), and genes encoding components of siderophore synthesis and transport (see Table 1 for details). To gain insight into the overall iron metabolism potential of *P. salmonis*, the bioinformatic tool FeGenie was used to identify genes coding for proteins involved in iron homeostasis. The results revealed that 23 genes related to iron transport, siderophore synthesis, siderophore transport, and gene regulation are present, 16 of them were up-regulated (log_2_ FC > 2) in the sgRNA-*fur1* strain (Table S4).

**Table 1.** Iron homeostasis genes upregulated in the knockdown strain. List of genes with significantly increased levels in the strain with induced expression of the silencing Mobile-CRISPRi system. Differentially expressed genes were grouped in three regions according to their location in *P. salmonis* genome.

| Gene ID | Log<sub>2</sub>FC | Gene name | Product description | Conserved domains | Predicted function |
| --- | --- | --- | --- | --- | --- |
| AWJ11_02085 | 3.52 | *feoC* | FeoC-like transcriptional regulator | Transcriptional regulator HTH-type | Fe<sup>2+</sup> transport |
| AWJ11_02090 | 3.13 | *feoB* | Transporter of a GTP-driven Fe<sup>2+</sup> uptake system | Ferrous iron transport protein B/ nucleoside | Fe<sup>2+</sup> transport |

|  |  |  |  | recognition |  |
| --- | --- | --- | --- | --- | --- |
| **AWJ11_02095** | 2.78 | *feoA* | Fe<sup>2+</sup> transport system protein A | FeoA domain | Fe<sup>2+</sup> transport |
| **AWJ11_06980** | 6.19 | *pvuE* | ABC transporter family protein | ABC-type cobalamin/Fe<sup>3+</sup>-siderophores transport system, ATPase component | Siderophore uptake |
| **AWJ11_06985** | 6.16 | *pvsE* | PLP-dependent decarboxylase | Pyridoxal-dependent decarboxylase, C-terminal sheet domain | Siderophore biosynthesis |
| **AWJ11_06990** | 8.63 | *pvsD* | IucA/IucC family protein | IucA / IucC family; ferric iron reductase FhuF-like transporter | Siderophore biosynthesis |
| **AWJ11_06995** | 8.33 | *pvsC* | MFS transporter | Arabinose efflux permease, multidrug efflux MFS transporter | Siderophore export |
| **AWJ11_07000** | 9.24 | *pvsD* | IucA/IucC family protein | IucA / IucC family; ferric iron reductase FhuF-like transporter | Siderophore biosynthesis |
| **AWJ11_07005** | 9.84 | *pvsA* | ATP-dependent carboxylate-amine ligase | ATP-grasp domain | Siderophore biosynthesis |
| **AWJ11_07010** | 10.26 | *pirA* | TonB-dependent siderophore receptor | TonB-dependent receptor plug domain; TonB dependent receptor-like, beta-barrel | Siderophore uptake |
| **AWJ11_07015** | 5.73 | *exbB* | Biopolymer transporter ExbB | MotA/TolQ/ExbB proton channel family | Siderophore uptake |
| **AWJ11_07020** | 5.99 | *exbD* | Biopolymer transporter ExbD | Biopolymer transport protein ExbD/TolR | Siderophore uptake |
| **AWJ11_07025** | 5.80 | *exbD* | Biopolymer transporter ExbD | Biopolymer transport protein ExbD/TolR | Siderophore uptake |
| **AWJ11_07030** | 7.78 | *tonB* | Energy transducer TonB | Gram-negative bacterial TonB protein C-terminal | Siderophore uptake |
| **AWJ11_07035** | 5.51 | *fepB* | Iron-siderophore ABC transporter substrate-binding protein | Periplasmic binding protein | Siderophore uptake |
| **AWJ11_07040** | 5.49 | *fepCD* | Iron ABC transporter permease | FecCD transport family | Siderophore uptake |
| **AWJ11_07380** | 7.24 | *bfd* | (2Fe-2S)-binding protein | BFD-like [2Fe-2S] binding domain | Iron mobilization |

In addition, small but significant changes in the expression of the down-regulated genes were detected in the IPTG-induced sgRNA-*fur1* strain compared to the uninduced strain, with only 24 genes, including the *fur* gene, showed a log_2_ fold change ≤ −1 (Table S4). Among them, we found genes encoding Fe-S-, heme- or iron-containing proteins such as, *sodB*, *thiC*, *cydB*, *nrdAB*, *acnA* (Table S4). Validation by qPCR of the expression profiles of representative genes up- or down-regulated in the IPTG- induced sgRNA-*fur1* strain confirmed the expression changes determined by the RNA-seq (Fig. 6C). In accordance with the increased expression of iron acquisition genes, TXRF analysis revealed a significant rise in intracellular iron concentration in the IPTG-induced sgRNA-fur1 strain relative to the non-induced strain (Fig. 6D).

### Bioinformatics prediction of Fur-binding sites in up-regulated genes

Using the *P. salmonis* CGR02 genome, we examined the intergenic regions upstream of up-regulated genes to identify sequences corresponding to the Fur (Fur box) binding site, based on experimentally validated sites from the CollectTF database (Kiliç *et al*., 2014). We found 16 potential binding sites within the 45 genes with log_2_ FC ≥ 1 in the IPTG-induced sgRNA-*fur1* strain (Table S4). Fifteen of these sites correspond to binding sites upstream of genes with log_2_ FC ≥ 2. Orphan Fur boxes were identified in two regions: 77 pb upstream of the gene *feoA,* in the FeO cluster, and 41 pb upstream a hypothetical protein with a methyltransferase domain gene (AWJ11_07530). In the remaining regions, there are clusters of Fur boxes, consisting of two to four sites. For example, four binding sites can be found upstream of genes from PirA-PvsACDE-PvuE cluster, and three sites were identified upstream of genes from members of the TonB-ExbB-ExbD energy transduction system. Also, single unit of transcription as the (2Fe-2S)-binding protein gene *bfd* and the gene AWJ11_07375 exhibit two and three Fur boxes, respectively (Fig. 7). We align the sequences of the 17 binding sites and identify a 9-1-9 bp inverted repeat sequence compatible with the classic Fur box model GATAATGATAATCATTATC described in *E. coli* (Escolar *et al*., 1999) or the two overlapping 7-1-7 TGATAAT sequence, described in *B. subtillis* (Fuangthong & Helmann, 2003).

**Fig. 7.**
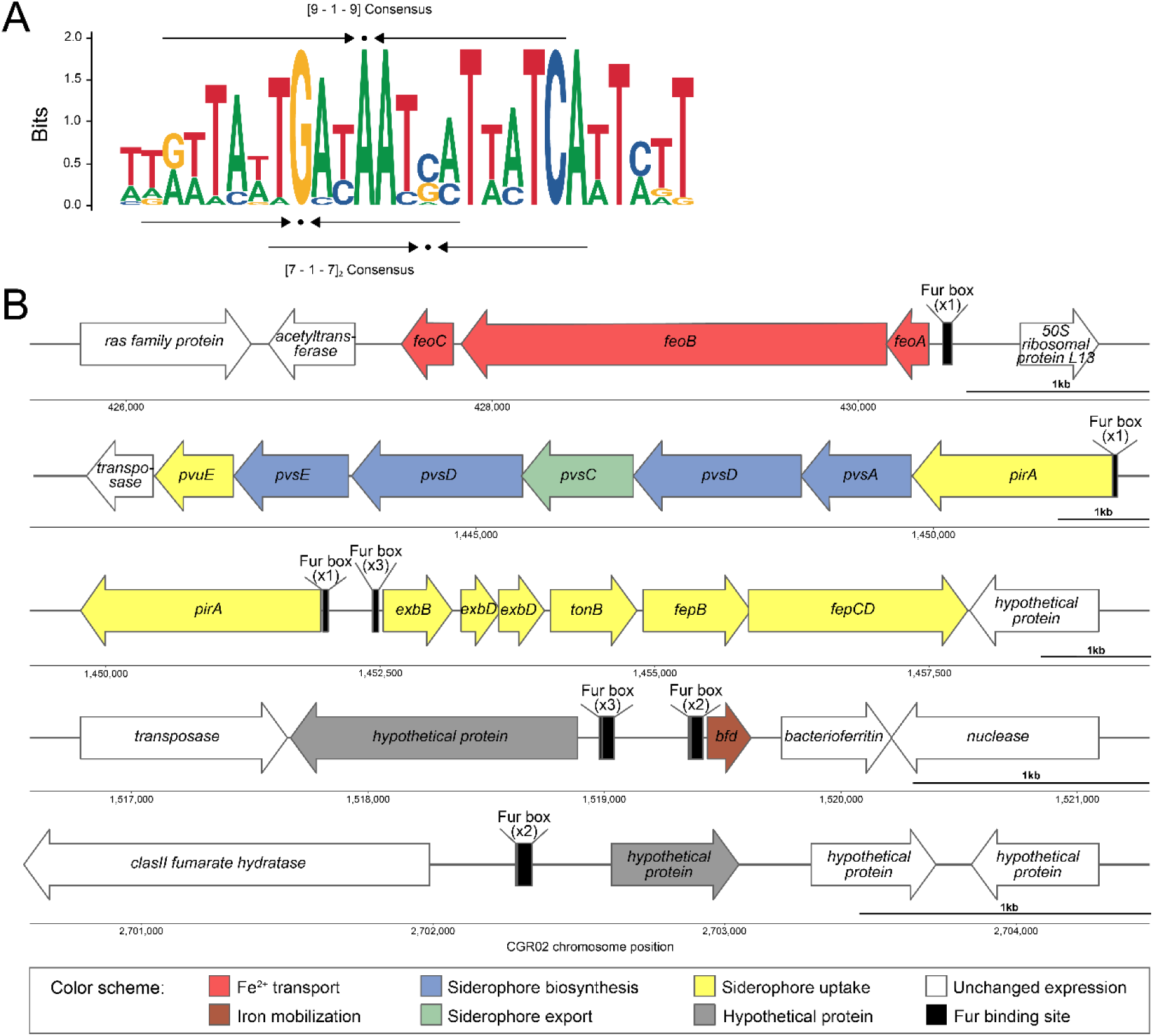
Fur-regulated genes in *P. salmonis* CGR02. (A) Logo of Fur box. The image shows the consensus logo for the Fur transcription factor binding sites identified upstream of genes that were upregulated by IPTG induction. The consensus fits the two-dimer model (9-1-9, top arrows) as well as the repeated heptamer model (7-1-7, bottom arrows) for fur binding. (B) Schematic representation of the up-regulated genes shown in Table 1. Predicted Fur boxes are indicated in black upstream of the regulated gene(s).

## Discussion

Implementation of Mobile-CRISPRi system in the fish pathogen *P. salmonis P. salmonis* is a major salmon pathogen with worldwide distribution and the principal cause of infection-related deaths in the Chilean salmon industry, having an important economic effect in the second largest exporter of salmon after Norway. Using genetic tools to study non-model organisms, such as the fastidious intracellular pathogen *P. salmonis* (Rozas-Serri, 2022), is essential to understanding gene function and furthering our knowledge of bacterial pathogenesis and disease. To date, only one gene manipulation technique has been proven to work for homologous recombination in *P. salmonis* (Mancilla *et al*., 2018); however, this approach is complex and time-consuming. Conversely, the CRISPRi gene knockdown editing system is a more advanced synthetic biology tool that is widely used in many microorganisms due to its simplicity and tunability, which can be achieved by optimizing the inducible expression of its components (Banta *et al*., 2020; Larson et al., 2013; Peters *et al*., 2019). Here, we took advantage of the Mobile-CRISPRi system described by Banta *et al*. (Banta *et al*., 2020) for the reversible and inducible knockout of target genes in *P. salmonis*, an exogenous *sfGFP* gene encoding for a fluorescent marker protein to test the functionality of the silencing system in this bacterium, and an endogenous *P. salmonis* gene to further investigate its function. The introduction of the inactivating element by conjugation of a suicide vector in *P. salmonis* resulted successful for inserting the Tn7 element in the *attTn7* box downstream of the *glmS* gene (as expected in Gamma-proteobacteria), which was highly conserved for the 92 *P. salmonis* genomes publicly available. We proved that the Mobile-CRISPRi system was expressed in the *P. salmonis* genetic context, and conferred gentamycin resistance and fluorescence to the receptor strain. We also observed basal *dcas9* expression levels in un-induced cells, and an increment of over 20- to 60-fold in expression after induction with IPTG. This increment did not impact on the growth of *P. salmonis* strains (Figure 3 and 5C). The leaky nature of *dcas9* expression has been reported previously (Rosano & Ceccarelli, 2014), but, in our experiments, this basal dCas9 level did not decrease the fluorescence levels as observed by comparing the fluorescence of the *P. salmonis* CGR02 *sfGFP*(+) strain (with constitutive expression of *sfGFP* in absence of the sgRNA) with the fluorescence from the *P. salmonis* CGR02 sgRNA-*sfGFP* strain (strain harboring the constitutively expressed *sfGFP* gene, and inducible *dcas9* and sgRNA) in the absence of the inducer (Figure 1E and 2B). The stability of Mobile-CRISPRi in the absence of selection was proved after over 130 generations (Figure 2), suggesting that this system could be a valuable tool for *in vivo* assays to study host-pathogen interactions, or other conditions where the presence of antibiotics is not suitable. Furthermore, not only was the Mobile-CRISPRi system inserted into the expected location and conferred the anticipated phenotypes to *P. salmonis*, but it also successfully silenced the expression of the target *sfGFP* gene in several *P. salmonis* strains from both genogroups, thus supporting its broad applicability. However, it is noteworthy that fluorescence intensity was not the same for the different strains, and strains from genogroup EM90-like (12201A and 8079A) and in addition to the Norwegian and Canadian strains (NVI 5692 and NVI 5892, respectively) presented a reduced fluorescence when compared to genogroup LF89-like strains (particularly CGR02 and PSCGR01) (Figure 4), thus suggesting that gene expression, protein translation and/or codon usage could differ between these strains.

### Silencing *fur* expression in *P. salmonis*

*P. salmonis* can utilize both ferric and ferrous iron, as well as synthesize siderophores, under iron-restricted conditions (Calquín *et al*., 2018). This suggests that the bacterium possesses various iron transport systems. In line with this, previous bioinformatics analysis has shown that the LF-89 strain of *P. salmonis* harbors a number of iron transport systems, as well as a gene that encodes a Fur homologue (Pulgar *et al*., 2015). This gene was subsequently characterized functionally through heterologous expression in a *Salmonella* Typhimurium Δ*fur* strain (Almarza *et al*., 2016). In this study, we examined the impact of inactivating *P. salmonis* Fur under conditions of moderately elevated intracellular iron concentration that did not affect bacterial viability using the Mobile-CRISPRi system in combination with transcriptomics. Our prior knowledge of the classical role of Fur in coordinating iron metabolism with iron availability, by regulating iron uptake and acquisition systems, made this study possible. Our results showed that, in the sgRNA-*fur1* strain, genes involved in iron acquisition are no longer regulated by iron and are highly expressed even under conditions of iron supplementation. This suggests that *P. salmonis* Fur acts as a repressor of these genes under conditions of high intracellular iron concentration.

In many bacteria, Fur negatively regulates the production and transport of siderophores (Troxell & Hassan, 2013) Consistent with this, upregulation of 13 genes predicted to be involved in siderophore synthesis and transport (Table 1) was detected in the sgRNA-fur1 strain. The high level of differential expression of genes encoding a TonB-dependent siderophore receptor and putative siderophore biosynthetic proteins in the knockdown strain is consistent with the prediction that Fe³⁺ predominates during *P. salmonis* infection cycles owing to iron sequestration by host proteins. In addition, previous studies have shown that *P. salmonis* can use ferric iron in both *in vitro* and *in vivo* infection models. This was evidenced by a decline in growth and enhanced resistance to *P. salmonis* infection following exposure to the ferric iron chelator deferoxamine (DFO, Díaz *et al*., 2021). Our transcriptomic data also suggests that *P. salmonis* has an additional method of iron acquisition: FeoB-mediated Fe²⁺ uptake. Previous work indicates that *P. salmonis* resides in a specialized phagosome within the host cell (Zúñiga *et al*., 2020). Thus, if the pH of the phagosome decreases, the amount of soluble ferrous iron may increase, making it available for uptake through the FeoAB system (Gammella *et al*., 2017). In this sense, *Legionella pneumophila*, which is phylogenetically related to *P. salmonis* (Bontemps *et al*., 2024), requires the function of FeoB when growing in low-iron conditions or inside the host cells (Cianciotto, 2015).

Upregulation of the *bfd* gene, which encodes a bacterioferritin-associated ferredoxin, was also detected. Bfd is required to mobilize iron stored in bacterioferritin (Yao *et al*., 2016), and its gene has been shown to be upregulated under iron-starved conditions in other bacterial pathogens (Avican *et al*., 2021; McHugh *et al*., 2003; Ochsner *et al*., 2002). In *Pseudomonas aeruginosa*, knocking out the *bfd* gene leads to an irreversible accumulation of iron in storage proteins and an iron deficiency in the cytosol (Eshelman *et al*., 2017). Furthermore, cell death has been observed in mature biofilms when inhibitors of the interaction between Bfd and bacterioferritin are used (Soldano *et al*., 2021). These findings emphasize the significance of this system in maintaining iron homeostasis and suggest a potential new therapeutic target for treating infections caused by pathogenic bacteria (Rivera, 2023).

It is known that iron-complexed Fur can also act indirectly in the regulation of some genes by repressing the expression of RyhB small RNA (Massé & Gottesman, 2002). The function of RyhB is to repress non-essential proteins that use iron as a way of preserving the iron levels in the cell at low iron conditions (Massé *et al*., 2005). Interestingly, among downregulated genes we identified genes encoding Fe-S-, heme- or iron-containing proteins such as, SodB, ThiC, CydB, NrdAB, AcnA, suggesting that these genes might be targets of RyhB. Indeed, the genes *sodB*, *cydB* and *acnA* are recognized targets of RyhB in *E. coli* (Massé *et al*., 2005). To date, two reports have predicted the presence of non-coding RNA molecules in the *P. salmonis* genome (Segovia *et al*., 2018). However, a RyhB homologue has not yet been identified. The absence of a RyhB homologue is likely to be attributable to two a low sequence similarity with known RyhB molecules. For instance, in *Pseudomonas aeruginosa* the small RNAs responsive to iron (PrrFs) share no sequence homology with *E. coli* RyhB (Wilderman *et al*., 2004). Overall, the transcriptional data were consistent with the expected role of Fur as a repressor of iron metabolism-associated genes in *P. salmonis*.

Our research also indicates that the regulation of overexpressed genes in *fur* knockdown occurs through previously characterized high-affinity sites for the Fur transcription factor (Lavrrar & McIntosh, 2003). The identification of such fur boxes, particularly within genes exhibiting more than twofold change in expression, strongly supports the role of Fur as a negative regulator. An analysis of the binding logo sequence indicates that the regulation of this gene can be achieved through a two-dimer of 5’-GATAAT-3’-5’ with each dimer overlapping on opposite sides of the DNA helix (F-F-x-R-R), comprising the 19-bp consensus binding site. As demonstrated by the observed regulation of ABC transporters within the enterobactin system, this is consistent with the primary source of iron uptake in *E. coli* (Lavrrar *et al*., 2002).

Beside gene clusters encoding iron acquisition and mobilization proteins, a number of other genes, mainly components of the type IV secretion system and hypothetical proteins, were also upregulated in the IPTG-induced sgRNA-*fur1* strain although with smaller changes in expression. Since consensus binding sites for Fur were not predicted within their intergenic regions, the observed differential expression might imply that other regulators may respond to iron concentrations within the cell. Possible examples of these regulators include Irr, and RirA (Small *et al*., 2009), neither of which are annotated in the genome of *P. salmonis*. However, a transcriptional regulator from the Rrf2 family to which RirA belongs has been predicted. Thus, it appears that other iron-responsive regulators may be present in *P. salmonis* but have not yet been identified.

Whilst there are a number of technical considerations to take into account when selecting a knockdown method, such as transcript stability and transcript and protein half-life, the Mobile-CRISPRi system described here can be used to efficiently and reversibly inhibit the expression of *P. salmonis* genes. This provides an alternative to traditional engineering approaches and a powerful tool to study gene function in this fastidious intracellular bacterium. We further expect it will enable new and fundamental experiments for host-pathogen interaction.

## Acknowledgments

This work was supported by grants from Agencia Nacional de Investigacion y Desarrollo de Chile (ANID), Fondecyt 1211893 (VC), Millennium Science Initiative Program: ICN2021_044 (VC, MG) and Doctoral fellowship 2024T2DID (PA). The authors thank Sebastian Roque for his assistance in the laboratory.

